# Comprehensive mapping and modelling of the rice regulome landscape unveils the regulatory architecture underlying complex traits

**DOI:** 10.1101/2024.06.24.600524

**Authors:** Tao Zhu, Chunjiao Xia, Ranran Yu, Xinkai Zhou, Xingbing Xu, Lin Wang, Zhanxiang Zong, Junjiao Yang, Yinmeng Liu, Luchang Ming, Yuxin You, Dijun Chen, Weibo Xie

## Abstract

Unraveling the regulatory mechanisms that govern complex traits is pivotal for advancing crop improvement. Here we present a comprehensive regulome atlas for rice (*Oryza sativa*), charting the chromatin accessibility across 23 distinct tissues from three representative varieties. Our study uncovers 117,176 unique open chromatin regions (OCRs), accounting for ∼15% of the rice genome, a notably higher proportion compared to previous reports in plants. Integrating RNA-seq data from matched tissues, we confidently predict 59,075 OCR-to-gene links, with enhancers constituting 69.54% of these associations, including many known enhancer-to-gene links. Leveraging this resource, we re-evaluate genome-wide association study results and discover a previously unknown function of *OsbZIP06* in seed germination, which we subsequently confirm through experimental validation. We optimize deep learning models to decode regulatory grammar, achieving robust modeling of tissue-specific chromatin accessibility. This approach allows to predict cross-variety regulatory dynamics from genomic sequences, shedding light on the genetic underpinnings of cis-regulatory divergence and morphological disparities between varieties. Overall, our study establishes a foundational resource for rice functional genomics and precision molecular breeding, providing valuable insights into regulatory mechanisms governing complex traits.

## Introduction

Rice *(Oryza sativa)* is not only one of the most important crops in the world but also an outstanding model species for studying plant growth and development. Over the past two decades, tremendous efforts have been made to understand the genetic basis of important agronomic traits in rice^1^. Genome-wide association studies (GWAS) have played a pivotal role in this pursuit, helping to link genetic variations to phenotypic diversity. These studies have identified a large number of candidate genes that hold promise for trait improvement^2-5^. However, despite these advances, our understanding of the regulatory mechanisms governing complex traits in rice remains incomplete.

Gene regulatory networks (GRNs) are largely dictated by *cis*-regulatory DNA sequences, such as promoters and enhancers, which are bound by specific transcription factors (TFs)^6^. Deciphering the regulatory code within these regulatory sequences and linking the regulatory sequences to target genes are crucial for rewiring GRNs for crop improvement and trait optimization^6^. Nonetheless, efforts to profile the regulome, encompassing the entirety of regulatory elements in the genome, remain constrained in rice. These efforts often concentrate on specific tissues, neglecting the comprehensive landscape across developmental stages and tissues^7,8^. Similarly, endeavors to establish links between regulatory regions and their target genes in rice are also limited^8^.

Meanwhile, many functional genetic variants associated with agronomic traits in rice reside within noncoding regulatory regions (e.g., *qSH1*^*9*^, *DROT1*^*10*^, and *FZP*^*11*^), which makes their interpretation challenging and underscores the necessity for a systematic dissection of regulatory sequences. Given that diverse traits manifest across distinct developmental stages and tissues, systematic annotation of noncoding regulatory variants in rice is currently hindered by the lack of a comprehensive epigenome map across various tissues and growth stages.

To bridge these gaps, we systematically mapped chromatin accessibility profiles in various tissues across the life cycle of three representative rice cultivars using the UMI-ATAC-seq method^12^, a modified ATAC-seq (assay for transposase accessible chromatin-sequencing) protocol developed in our lab. Through analysis of 145 ATAC-seq datasets, we obtained a total of 117,176 unique open chromatin regions (OCRs), accounting for *∼*15% of the rice genome. By integration of RNA-seq data from matched tissues, we predicted potential target genes for OCRs based on the correlation of gene expression and adjacent chromatin accessibility across tissues. Through TF footprinting analysis, we inferred tissue- or stage-specific regulatory networks and identified cultivar-polymorphic/trait-associated OCRs by comparing the regulatory landscapes between *indica* and *japonica* rice subspecies. Notably, our analysis unveiled a preference for GWAS-associated variants within tissue-specific OCRs, enabling the identification of causal associations between 209 complex agronomic traits and noncoding regulatory variants using this OCR landscape. Utilizing optimized deep learning models, we decoded the regulatory grammar through modeling of tissue-specific chromatin accessibility and across-variety predictions from sequences. The modeling approach sheds light on the key genetic alterations contributing to *cis*-regulatory divergence. Overall, these data not only serve as a cornerstone resource for the plant research community but also provide valuable regulatory variants for precision molecular breeding.

## Results

### Charting a reference atlas of chromatin accessibility in rice

To generate a comprehensive landscape of accessible chromatin in rice (*Oryza sativa*), we took advantage of an improved ATAC-seq protocol (UMI-ATAC-seq^12^, which incorporates unique molecular identifiers to the regular ATAC-seq technique for accurate quantification and footprinting) to perform chromatin accessibility profiling in 23 tissues/organs spanning the entire life cycle of rice. The representative tissues include callus, radicle, plumule, leaf, leaf sheath, root, apical meristem (AM1/AM2), dormant buds (DBuds), shoot apical meristem (SAM1/SAM2/SAM3), panicle neck node (PNN), stem, young panicle (Panicle1/Panicle2/Panicle3/Panicle4), lemma, palea, pistil, stamen and seed coat (Seed1/Seed2/Seed3). The experiments were conducted in three representative rice varieties, namely Nipponbare (NIP; *japonica* subspecies), Minghui 63 (MH63; *indica* subspecies type II), and Zhenshan 97 (ZS97; *indica* subspecies type I), with each experiment consisting of at least two biological replicates (Fig. 1a and Supplementary Data 1). In total, 145 genome-wide chromatin accessibility datasets with high sequencing depth (∼30.7M reads on average) were generated. We applied the ENCODE standards^13,14^ to establish the analysis pipeline (see *Methods*). Compared to published ATAC-seq datasets in the plants as deposited in the ChIP-Hub database^14^, our data exhibited a significantly higher signal-to-noise ratio (Supplementary Fig. 1). Through data analysis using the corresponding reference genomes of the three cultivars^15,16^, we identified on average of 40,676 (ranging from 28,991 to 49,737) reproducible OCRs (with an Irreproducible Discovery Rate [IDR]^17^ < 0.05) per experiment (Fig. 1b). As expected, the identified OCRs from all experiments predominantly located either in the proximal upstream regions of the transcription start site (TSS) or the distal intergenic regions (Fig. 1b,f, Supplementary Fig. 2 and Supplementary Data 2), resembling promoters or enhancers, respectively^18^. Of note, OCRs from intragenic regions accounted for a relatively small proportion (about 15.7%), while most of these OCRs originated from intronic regions (Supplementary Fig. 2b). These observations indicate that the vast majority of OCRs originate from noncoding regions of the rice genome.

**Fig. 1.**
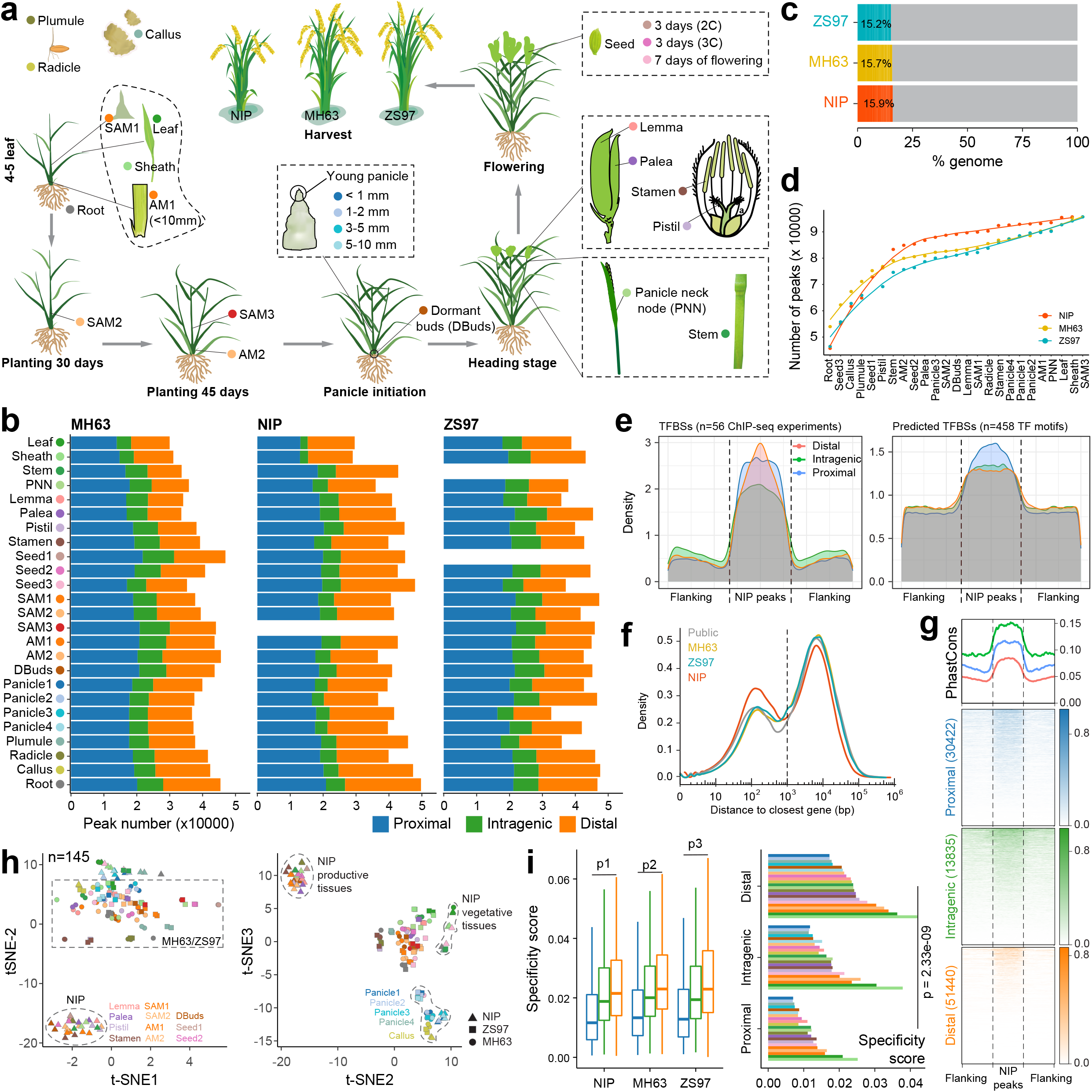
Characterization of an open chromatin landscape in rice. **a**. ATAC-seq and RNA-seq experiments were conducted in three varieties (Nipponbare, Minghui 63, and Zhenshan 97) of rice in various tissues across the entire life. See Supplementary Data 1 for detailed descriptions of sample collection. Consistent tissue color code is used throughout the figure **b**. Bar plot showing the number of reproducible OCRs identified from each tissue in the three rice varieties. The OCRs are classified into three categories based on the distance of the OCR summit to its closest transcription start site (TSS): distal (>1 kb), proximal (<= 1kb), and intragenic. No data from the tissues of SAM3 (NIP), Seed1 (ZS97) and Stem (ZS97). **c**.The proportion of the rice genome annotated as open chromatin regions (OCRs) in our study. **d**.The accumulative number of unique OCRs in each tissue, calculated by excluding OCRs that overlap with the OCR superset. **e**.Density plot showing the enrichment of TF binding sites (TFBSs) around the OCRs in Nipponbare (NIP). TFBSs were predicted either by ChIP-seq datasets for 56 distinct TFs (left) or DNA motifs for 458 TFs (right), which were obtained from the ChIP-Hub database ^14.^ The flanking area on both sides is 1kb. **f**.. The distribution of the distance of OCR summit to its closest TSS in the three rice varieties. Publised open chromatin data^14^ in rice (NIP) were included for comparison. Based on the distribution, a cutoff of 1 kb (dashed line) was used to distinguish the proximal and distal regulatory OCRs. **g**.The distribution of the conservation PhastCons score^19^ around the NIP OCRs. **h**.The t-SNE plot showing an unsupervised clustering analysis of chromatin accessibility across different samples. Each dot represents one replicate. Color code as in (a). **i**.Boxplot showing the distribution of tissue specificity score of intragenic (n= 14239), proximal (n=29524) and distal (n= 57153) OCRs (left) or the median score in each tissue. P1 = 4.01e-39, p2 = 2.13e-96, p3 = 1.23e-95. All *p*-values were calculated by two-sided Mann–Whitney U test between proximal and distal OCRs in terms of specificity. Tissue color code as in (a). Boxplot shows the median (horizontal line), second to third quartiles (box), and Tukey-style whiskers (beyond the box). Source data are provided as a Source Data file.

**Fig. 2.**
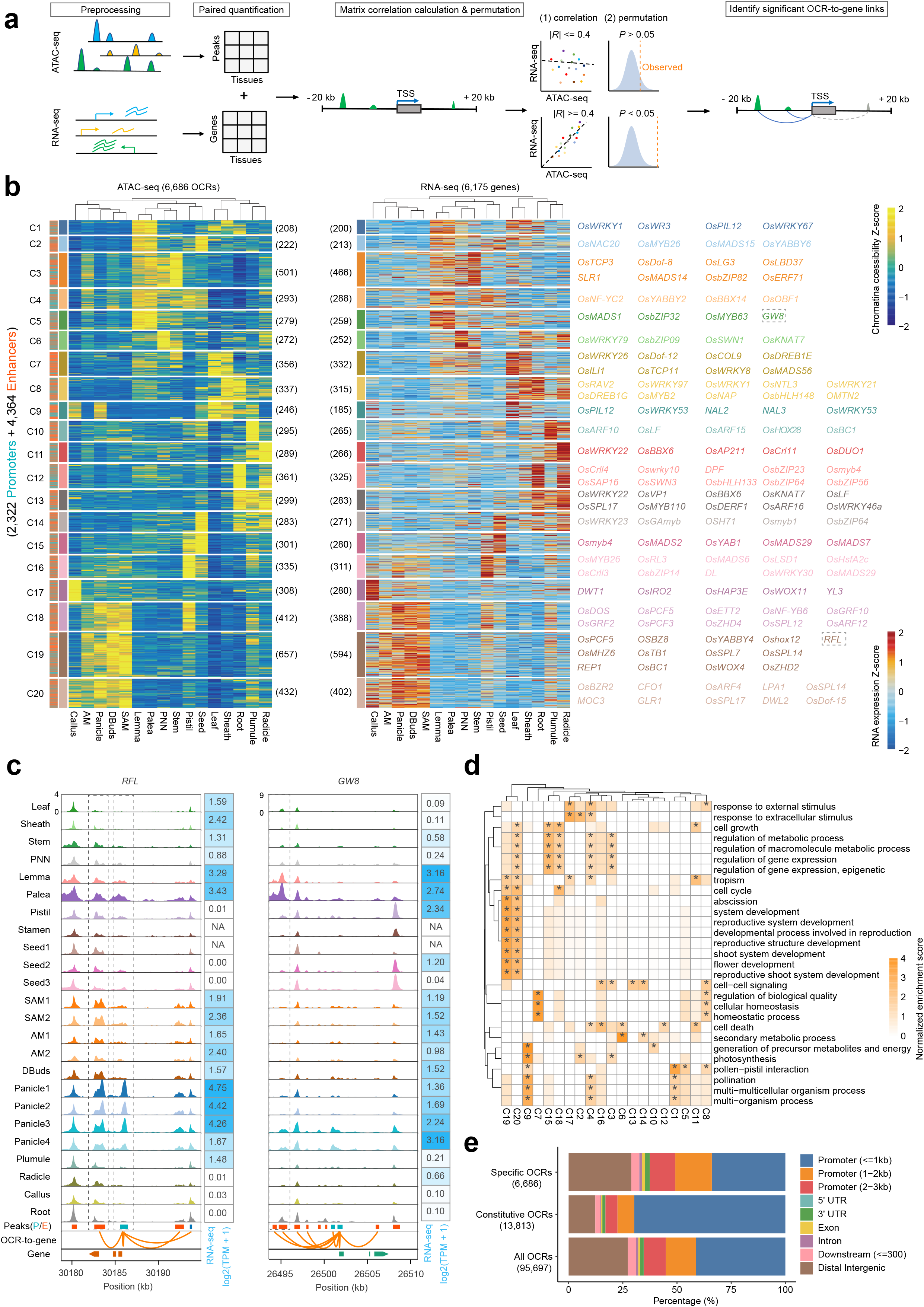
Tissue-specific OCRs. **a**.Schematic diagram illustrating the correlation-based approach to link ATAC-seq OCRs to target genes based on correlation analysis between chromatin accessibility and gene expression. **b**.Heatmap showing the tissue-specific OCR-to-gene links (*R* >= 0.4, *P* < 0.05, two-tailed *Z*-test). Each row in the left panel is a unique OCR. Each row in the middle panel is a gene, corresponding to target genes for OCRs in the left panel. Representative genes are shown on the right. **c**.Examples of tissue-specific OCRs (in the dashed box) regulating dynamic expression of the corresponding target genes. The orange lines indicate the OCR-to-gene links, and the deeper the line the higher the correlation between the chromatin accessibility and gene expression. **d**.Enrichment of biological processes gene ontology (GO) terms for target genes in each OCR cluster in (b). The asterisk (*) denotes *P* < 0.05 (*P*-values were calculated by Hypergeometric test after Bonferroni correction). **e**.Bar plot showing the percentage of OCRs from different categories based on the genomic location. Source data are provided as a Source Data file.

We estimated that approximate 15% of the rice genome could be annotated as OCRs, with a consistent pattern observed across each variety (Fig. 1c), and the estimation appeared to have reached saturation in rice (Fig. 1d). OCRs contain multiple TF binding sites and are responsible for regulating the expression of target genes^6,18^. We collected publicly available ChIP-seq data for 56 distinct TFs (Supplementary Data 3) and predicted DNA motifs for 458 TFs in rice from the ChIP-Hub database^14^, and showed that OCRs were significantly enriched for TF binding sites (Fig. 1e). Furthermore, we found that OCRs are highly evolutionarily constrained compared to flanking genomic regions (Fig. 1g), supporting previous findings that conserved noncoding sequences (CNSs) are predictive of OCRs in plants^19,20^.

We next assessed the overall similarities and differences of chromatin accessibility across varieties and tissues. We quantified all datasets based on the merged OCRs (n = 117,176) called from the same reference genome (i.e., Nipponbare) and visualized their global patterns using t-distributed stochastic neighbor embedding (t-SNE)^21^. While dimension 1 and 2 of t-SNE results generally reflected differences between the *indica* (MH63 and ZS97) and *japonica* (NIP) subspecies, dimension 2 and 3 primarily delineated distinct clusters among tissue types (Fig. 1h). For instance, the chromatin accessibility patterns of vegetative and productive tissues of NIP were separated into distinct clusters, whereas young panicles and callus tissues exhibited similar patterns regardless of their variety origin. We further calculated the tissue specificity of each OCR based on the Jensen-Shannon divergence (JSD) index. Obviously, distal OCRs showed significantly higher specificity scores than proximal OCRs (Fig. 1i and Supplementary Fig. 3a,b), consistent with previous findings^14,18,22^.

**Fig. 3.**
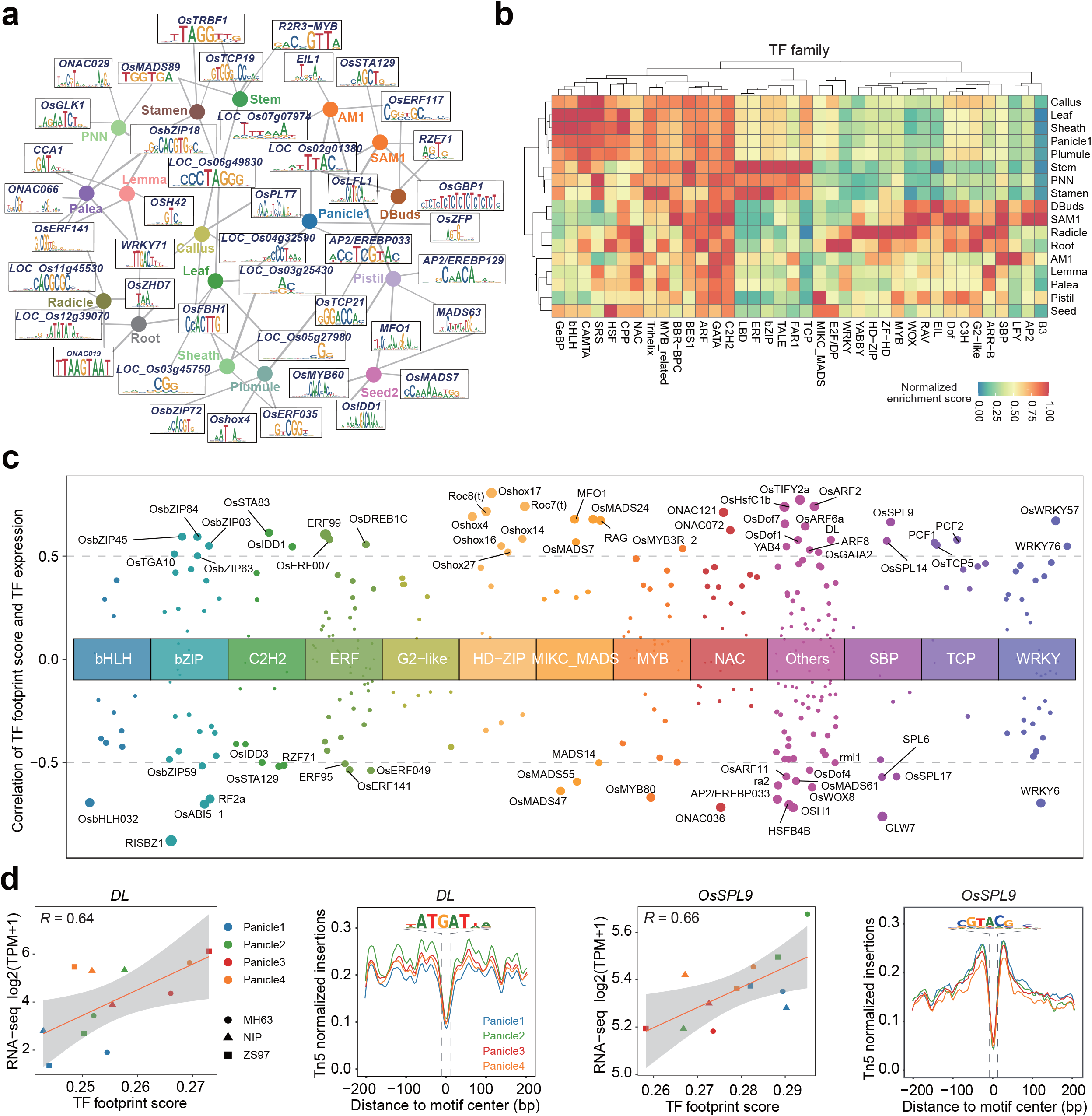
Tissues-specific and stage-specific regulatory elements. **a**.Enrichment of TF motifs in tissue-specific OCRs. Only top 5 enriched TFs in each tissue are shown. See Supplementary Data 8 for the full list. The thickness of edges is proportion to the corresponding enrichment score. **b**.The relative preference of regulators within TF families in each tissue type. Only the top 100 TF motifs in each tissue were used for analysis. **c**.The scatter plot showing the distribution of the Pearson correlation coefficient between TF footprint score and its expression. Only absolute values of correlation coefficients greater than 0.5 are marked. **d**. The scatter plot showing the distribution of TF footprint score and its gene expression in NIP, MH63, and ZS97(left). The error bands indicate 95% confidence intervals. Distribution of Tn5 cuts around the footprint of DL and OsSPL9 at different stages of young panicle (right). Source data are provided as a Source Data file.

In short, the comprehensive accessible chromatin landscape in rice represents a value resource for crop functional genomic studies.

### Linking open chromatin regions to target genes

To decipher which genes these OCRs may regulate, we generated matched RNA-seq datasets for the investigated tissues in each rice variety (Supplementary Fig. 3c and Supplementary Data 4). We adopted a strategy^23^ to predict OCR-to-gene links based on correlation analysis between the OCR accessibility and gene expression across all samples (Fig. 2a; see *Methods*). Genes can be regulated by multiple OCRs (including promoters and enhancers) through chromatin interactions, which are supposed to occur within topologically associated domains (TADs). Since the size of TADs in the rice genome was estimated to be ranging from 35 kilobase pair (kb) to 45 kb based on Hi-C data^24,25^, we restricted our analysis to 40 kb (i.e., from 20 kb upstream to 20 kb downstream of the TSS) to predict target genes of OCRs. Using a cutoff of absolute Pearson correlation coefficient |*R*| >= 0.4 and *P* < 0.05, we obtained a total of 59,075 unique links between OCRs (n = 38,437, 32.8% of all OCRs) and genes (n = 18,781, 48.1% of annotated genes; Supplementary Fig. 4a, b and Supplementary Data 5). As expected, the OCR-to-gene links tended to occur more frequently in the proximal OCRs, and consequently the correlation between the gene expression and chromatin accessibility is higher for proximal links (Supplementary Fig. 4c-f).

**Fig. 4.**
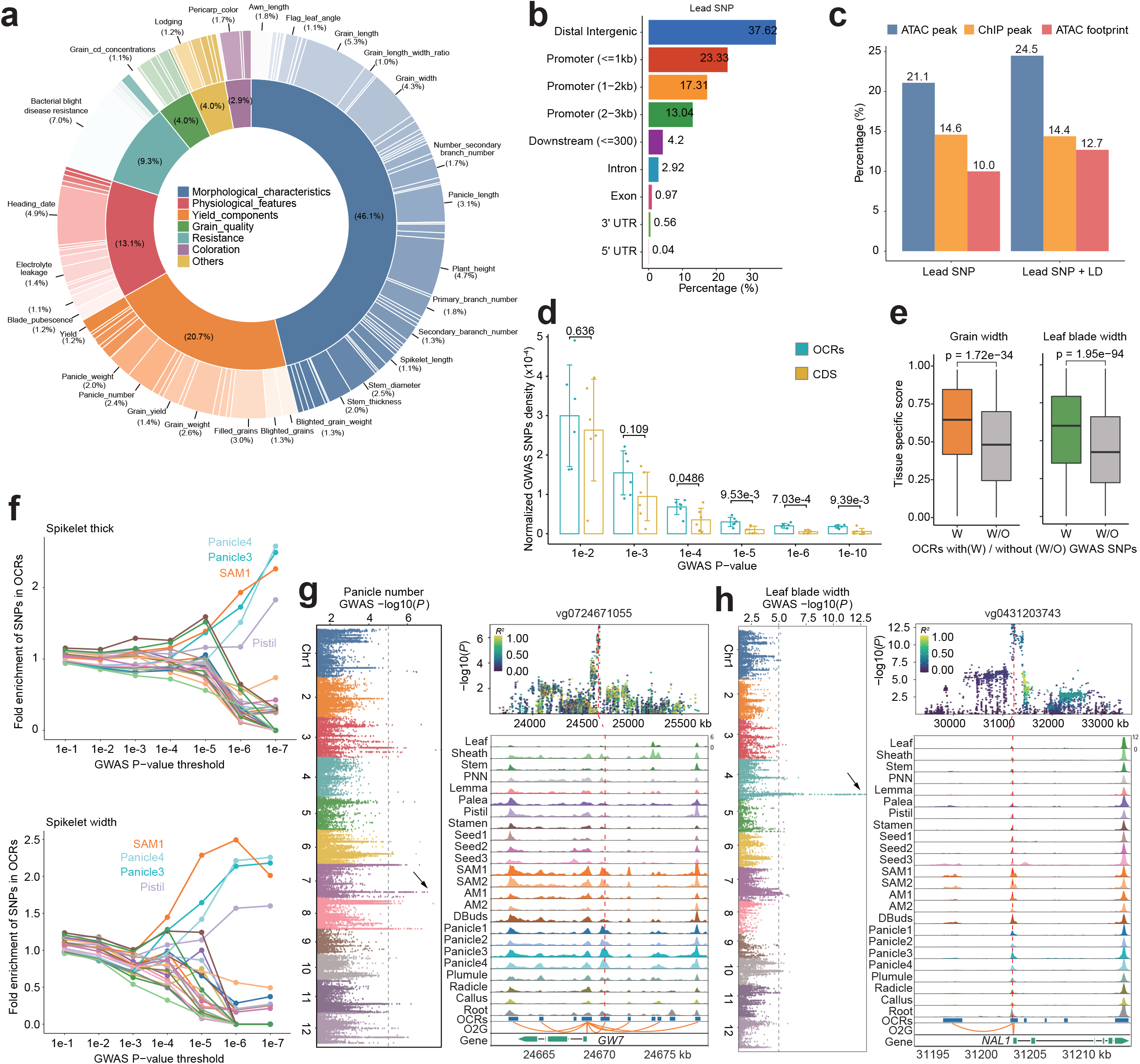
GWAS-associated variants localize in tissue-specific OCRs. **a**.Categorical proportions of lead SNP in each GWAS. The inner circle indicates the proportions of the seven major categories, and the outer circle indicates the subcategories contained in each major category. Only high proportions are marked in the outer circle. **b**.Distribution of curated lead SNPs by genomic context. All lead SNPs are the same as in (a). **c**.Overlap proportions of lead SNPs and sets of SNPs with strong linkage disequilibrium (LD > 0.8) with lead SNPs with ATAC-seq OCRs, ChIP-seq peaks and footprints identified by NIP ATAC-seq, respectively. **d**.The barplot showing the SNP density of OCR and CDS regions at different GWAS *P*-value thresholds. The error bars are the standard deviations of the SNP densities in the six GWAS catalogs from the (a). Data represents the mean⁳±⁳SD of 6 independent GWAS catalogs. The *P* values were calculated by two-tailed Student’s *t*-test. **e**.Boxplots showing the tissue-specificity score distribution of OCRs that overlap with grain width^54^ and leaf blade width^53^ GWAS SNPs. For grain width, the sample sizes for the “with” and “without” groups are 896 and 4480, respectively. For leaf blade width, the sample sizes for the “with” and “without” groups are 2864 and 5728, respectively. Boxplot shows the median (horizontal line), second to third quartiles (box), and Tukey-style whiskers (beyond the box). The *P*-values were calculated by two-tailed Student’s t-test. **f**.The enrichment of GWAS SNPs^2^ in OCRs with different GWAS *P*-value threshold. **g**. Manhattan plot showing the GWAS signal distribution of vg0724670482 and the LD distribution of its surrounding SNPs. The track plot demonstrates that the OCR where this SNP is located has a higher accessibility in palea tissue. “O2G” represents OCR-to-gene links. **h**. Same meaning as (g), except that vg0431203743 has a higher accessibility in SAM and young panicle. Source data are provided as a Source Data file.

Genetic variants within OCRs can contribute to changes in gene expression levels through expression quantitative trait loci (eQTL). We colocalized the identified OCR-to-gene links from our study with published eQTL data in rice^26^, and we found a significant overlap (Chi-squared test, *P* < 1.55e™06) between OCR-to-gene links and eQTL-gene pairs (Supplementary Fig. 4g). As expected, the correlation coefficients of colocalized OCR-to-gene links with eQTLs are significantly higher than those without colocalization (Wilcoxon test, *s* = 4.11e-38; Supplementary Fig. 4h). We identified numerous known regulatory variants that influence the expression of genes associated with agronomic traits. To name a few, a variant within a distal regulatory region (∼12 kb upstream) of *qSH1* modulates its expression dynamics, leading to change the seed shattering in rice^27^. Accordingly, there is a positive correlation (*R* = 0.47, *P* < 0.013) between the accessibility of this enhancer and the expression of *qSH1* in various tissues, particularly in SAM where gene expression increases (Supplementary Fig. 4i,l). Similarly, *OsLG1* is tightly linked to upstream regulatory regions that colocalize with a strong QTL associated with the panicle shape trait^28^ (Supplementary Fig. 4j,l). *IPA1* showed significantly positive correlation (*R* = 0.84, *P* < 2.95e-8) between its enhancer activity and gene expression, with increased expression in yield-related tissues (Supplementary Fig. 4k,l), confirming an important role of *IPA1* to shape rice ideal plant architecture (IPA) and thus to enhance grain yield^29^.

Taken together, the predicted OCR-to-gene links provide regulatory insights into agronomic trait development in rice and highlight targetable OCRs of important genes for genome editing.

### Dissecting tissue-specific and stage-specific regulatory grammar

The comprehensive chromatin accessibility landscape of representative tissues gave us an opportunity to uncover tissue-specific regulatory grammar. We quantified the tissue-specificity of OCRs by utilizing the JSD score, which enables the discrimination of target genes from housekeeping (e.g., *GAPDH*^*30*^ and *OsGOGAT1*^*31*^) to tissue-specific (e.g., *OsYABBY5*^32^ and *OsWRKY47*^*33*^) according to the above predicted OCR-to-gene links (Supplementary Fig. 5 and Supplementary Data 6). We have specifically focused on analyzing highly tissue-specific OCRs (n = 6,686 with a cutoff of JSD > 0.08, ∼ 7% of all OCRs) as they may encode the tissue-specific regulatory grammar. These OCRs were further annotated as promoters (n = 2,322) or enhancers (n = 4,364) according to the genomic distance to the TSS. By performing joint clustering analysis of chromatin accessibility and target gene expression using OCR-to-gene links, we identified 20 distinct clusters of OCRs (Fig. 2b and Supplementary Data 7). Each cluster had 200∼500 OCR-to-gene links that were highly activated in specific tissues, and showed a high degree of consistency with the known biological characteristics of the corresponding tissues (Fig. 2b-d). For instance, the palea- and lemma-specific links in cluster 5 (C5) contained promoter-enhancer interactions at the locus of *GW8*, which is a known gene controlling grain weight in rice^34^ (Fig. 2c). Accordingly, *GW8* was highly expressed in pistil, lemma, and palea. Gene ontology (GO) enrichment analysis using genes from C5 revealed that biological processes such as ‘pollen™pistil interaction’ and ‘pollination’ were overrepresented (Fig. 2d). Similarly, we identified a number of OCRs in C19 that were highly and specifically accessible in meristem-like tissues (including young panicle and shoot apical meristem), and the associated target genes showed significant enrichment for functions related to ‘reproductive system development’, ‘flower development’, and ‘shoot system development’ (Fig. 2b, d). Notably, *RFL*, a crucial regulator for plant architecture and flowering time^35,36^, was among these target genes (Fig. 2c). Interestingly, we observed that a higher proportion (28.9%) of tissue-specific OCRs originated from distal intergenic regions compared to constitutive OCRs (12.3%). In contrast, approximately 85% of constitutive OCRs were derived from the proximal-promoter regions. (Fig. 2e).

**Fig. 5.**
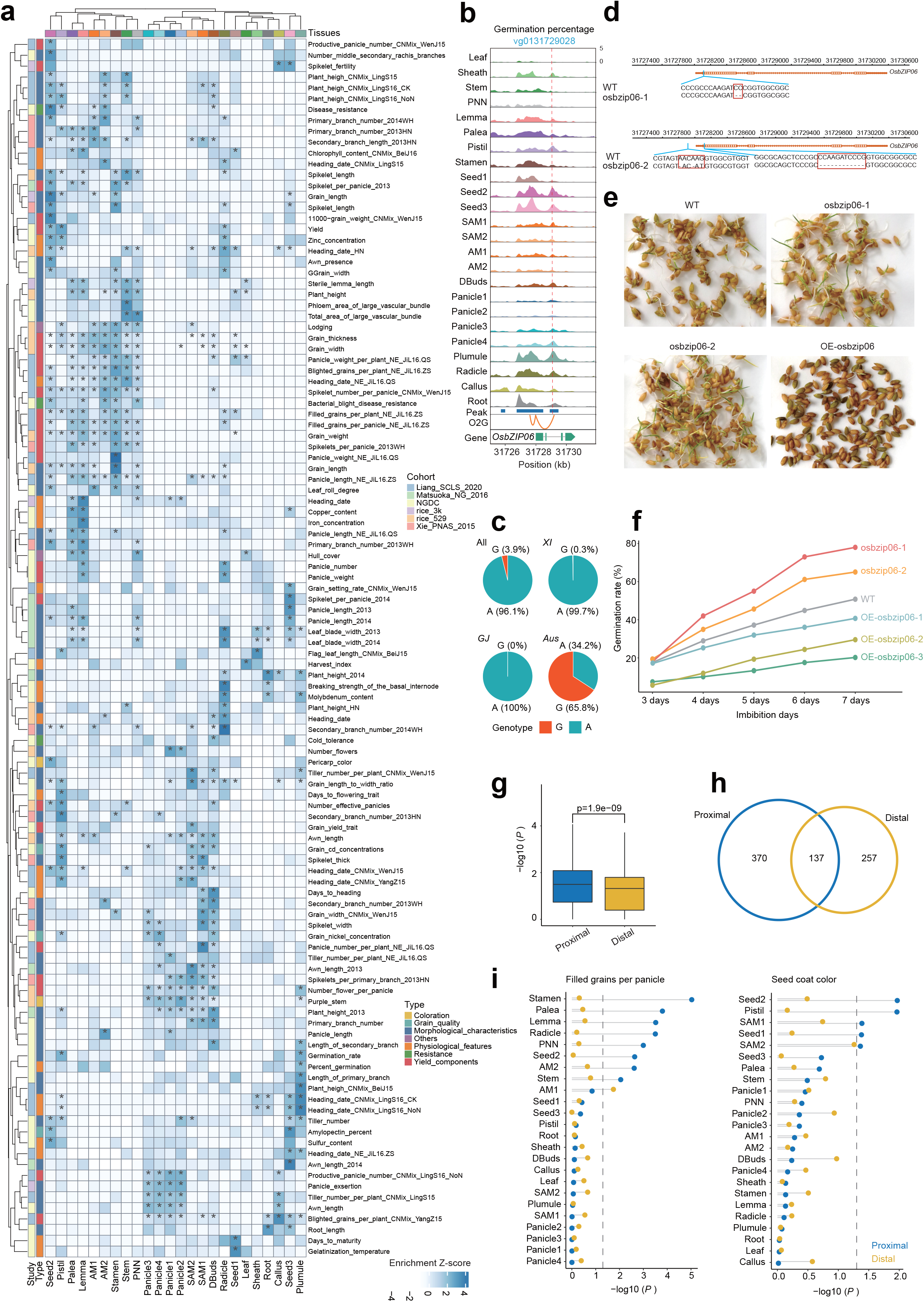
Association of tissue type with complex traits. **a**.GWAS SNPs enrichments for ATAC-seq OCRs of different tissues. The heatmap showing the significant tissue-specific enrichment results. The values are transformed by -log10(P) and then normalized by row. Those marked with an asterisk represent *P* < 0.05 for this result. The *P* values were calculated by Kolmogorov-Smirnov test. Only tissue data for the NIP variety were used for this analysis. The full list for GWAS enrichment result could access by Supplementary Data 11. **b**.One representative examples of genomic tracks at loci *OsbZIP06* showing that GWAS lead SNP is located in tissue-specific OCRs. The GWAS study name and SNP location (denoted by red dashed line) are shown at the top of panel. **c**.Haplotype distribution of vg0131729028 in the population. This result was obtained from the RiceVarMap 2.0 database^7^. **d**.Identification of mutation information of two *OsbZIP06* mutants based on sanger sequencing. **e**. The images show seed germination rates of wild type and mutants of *OsbZIP06*. **f**.The line graph showing the germination rates of different mutants *osbzip06* at different days of imbibition. “OE” represents overexpression. **g**.Boxplot showing the enrichment results of proximal and distal OCRs with 209 GWAS results respectively. Only results where GWAS was significantly enriched with at least one of proximal and distal OCRs are shown. The sample size of each group is 764. The *P* value was calculated by Student’s t-test. Boxplot shows the median (horizontal line), second to third quartiles (box), and Tukey-style whiskers (beyond the box). **h**.Venn plot showing the number of results significantly enriched (*P* < 0.05, Kolmogorov-Smirnov test) by proximal and distal OCRs. **i**.Enrichment of GWAS SNPs in TSS proximal and distal OCRs. The names of the GWAS are marked at the top of the panel. The grey dashed line indicates the *P*-value threshold of 0.05. The *P* values were calculated by Kolmogorov-Smirnov test. Source data are provided as a Source Data file.

To delineate the TFs that may bind to these tissue-specific OCRs, we used GimmeMotifs^37^, a versatile tool can detect tissue-specific TF binding motifs by comparing TF binding activity across multiple experiments. We restricted our analysis to the top 2,500 OCRs in each tissue, as determined by their specificity measurement (SPM) score^38^. The predicted regulatory motifs showed significant enrichments in a tissue-specific manner in matching tissue types (Supplementary Fig 6 and Supplementary Data 8). We narrowed our focus to the top enriched regulators in each tissue type, and found many of the inferred links correspond to known regulatory relationships (Fig. 3a). For example, *OsIDS1*, a gene that plays a vital role in shaping inflorescence structure and establishing floral meristems^39,40^, exhibited relatively high activity in the panicle. *OsbZIP72*, enriched in plumule tissue, has been found to regulate plumule length by modulating abscisic acid (ABA) signaling and promote seed germination^41,42^. Notably, the tissues of seed and pistil demonstrated a co-enrichment pattern of crucial regulators involved in flower and seed development, including *MFO1* and *MADS63*^*43-45*^ from the MADS gene family (Fig. 3a). For each tissue type, we performed a systematic analysis to calculate the relative preference of regulators within TF families. Our analysis revealed distinct tissue-specific TF binding patterns, indicating clear preferences for specific regulators in different tissues (Fig. 3b). For instance, the TCP TF family showed a preference for enrichment in stem, stamen, and panicle neck node (PNN) tissues. This observation aligns with the known biological function of TCP genes, specifically their role in regulating cell proliferation in developing tissues^46^.

**Fig. 6.**
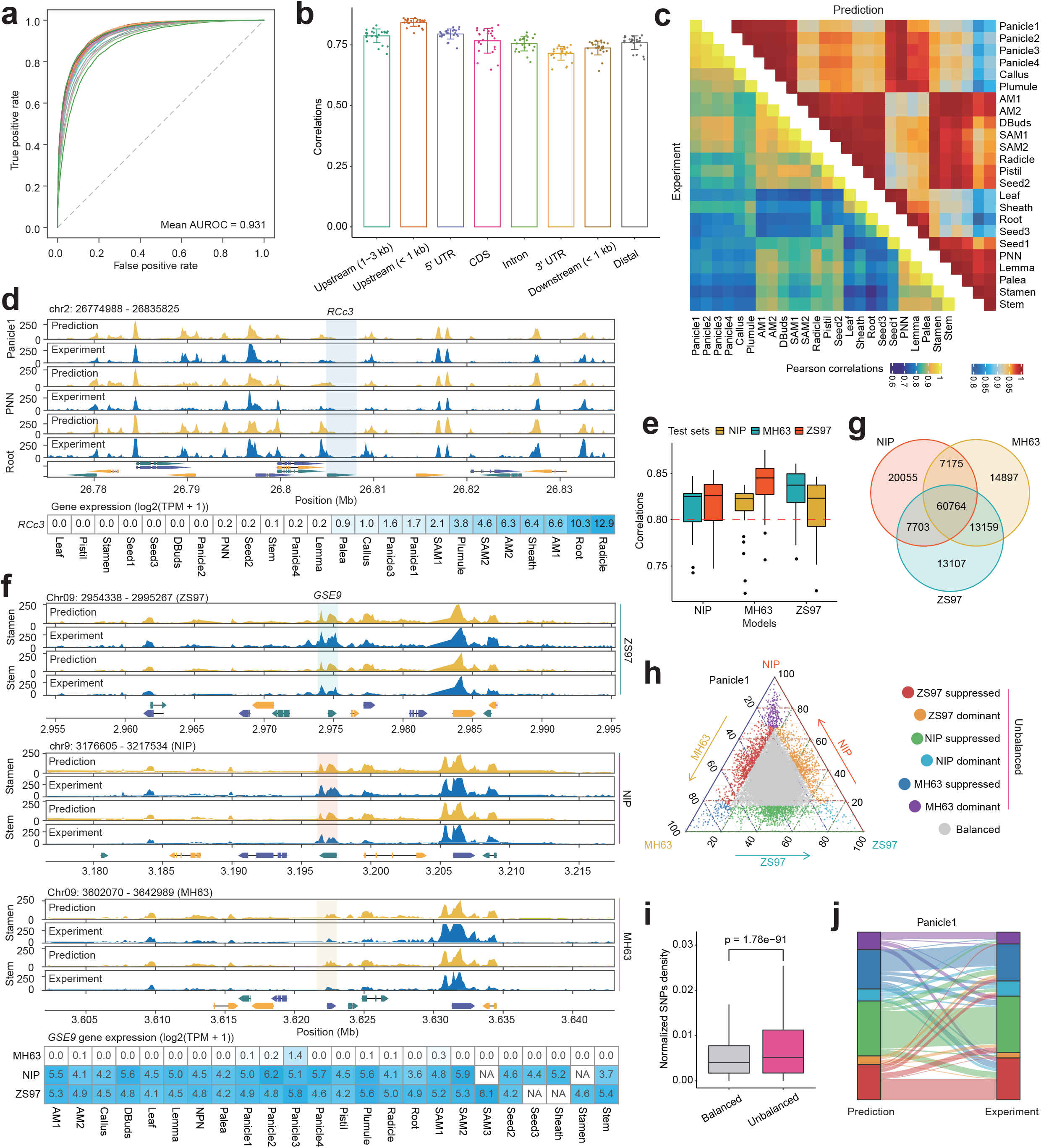
Using deep learning model to predict chromatin accessibility across tissues and varieties. **a**.Receiver operating characteristic curves for different tissues in the NIP cultivar. The average AUORC value was 0.931. **b**.Distribution of Pearson correlation coefficients between predicted and true signal values for different genomic regions using NIP model. Each point represents one tissue (n = 24). Data are displayed as mean ± SD. **c**.Comparison of clustering results based on predicted and true signal values using NIP model. **d**.The genomic tracks show the signal values predicted by NIP model versus the true signal values for Panicle1, PNN and Root, respectively. The shaded area is labelled with the gene region of RCc3. The heatmap below the tracks show the expression of the RCc3 in NIP varieties. **e**.The boxplot showing the distribution of Pearson’s correlation coefficients for the models of NIP, MH63 and ZS97 tested separately using sequences from the other two varieties. The red dashed line represents a correlation coefficient at 0.80. Each sample consists of 24 observations. Boxplot shows the median (horizontal line), second to third quartiles (box), and Tukey-style whiskers (beyond the box). **f**.The genomic tracks showing the signal values predicted with the ZS97 model for NIP, MH63 and ZS97 sequences versus the true signal values in Stamen and Stem tissues, respectively. The shaded area represents the orthologous region of *GSE9* in NIP, MH63 and ZS97 varieties. The heatmap below the tracks show the expression of the *GSE9* in NIP, MH63, and ZS97 varieties. **g**.Comparison of OCRs in the three rice cultivars (NIP, MH63 and ZS97). For each cultivar, OCRs from all tissues were merged and then compared based on whole genome sequence alignments. **h**.Ternary plot showing the chromatin accessibility of orthologous OCRs among the three rice cultivars with Panicle1 tissue. **i**.Comparison of the SNP density within the balanced (n=19793) and unbalanced (n=8385) orthologous OCRs. The *P* value was calculated by two-tailed Student’s *t* test. Boxplot shows the median (horizontal line), second to third quartiles (box), and Tukey-style whiskers (beyond the box). **j**.Sankey diagram showing the true chromatin accessibility difference and the chromatin accessibility difference predicted by the deep learning model for orthologous OCRs in NIP, MH63 and ZS97. The color representation is categorized in the same way as in (**h**). Source data are provided as a Source Data file.

Analyzing temporal ATAC-seq data through footprinting could assist in identifying key regulators, such as pioneer factors, that control developmental progression and transition^47^. We generated temporal open chromatin data from the young panicle, which is a crucial organ determining the yield of rice^48,49^, across four successive developmental stages (<1 mm, 1-2 mm, 3-5 mm, and 5-10 mm; Fig. 1a). We endeavored to identify regulatory motifs that exhibited either positive or negative correlation with the young panicle developmental stage in terms of enrichment, using dynamically changing OCRs (n = 9,244; Fig. 3c, Supplementary Fig. 7a and Supplementary Data 9). The regulators that were most enriched displayed predominantly positive correlations, indicating their function as transcriptional activators. Conversely, a subset of factors exhibited negative correlations, suggesting a repressive role. In this regard, DL (encoding OsYABBY^50^), OsSPL9^51^ and OsSPL14^52^ were identified as representative positive regulators, during the development of young panicles in rice (Fig. 3d and Supplementary Fig. 7b). However, further experimental data is necessary to validate the potential involvement of these TFs in young panicle development.

**Fig. 7.**
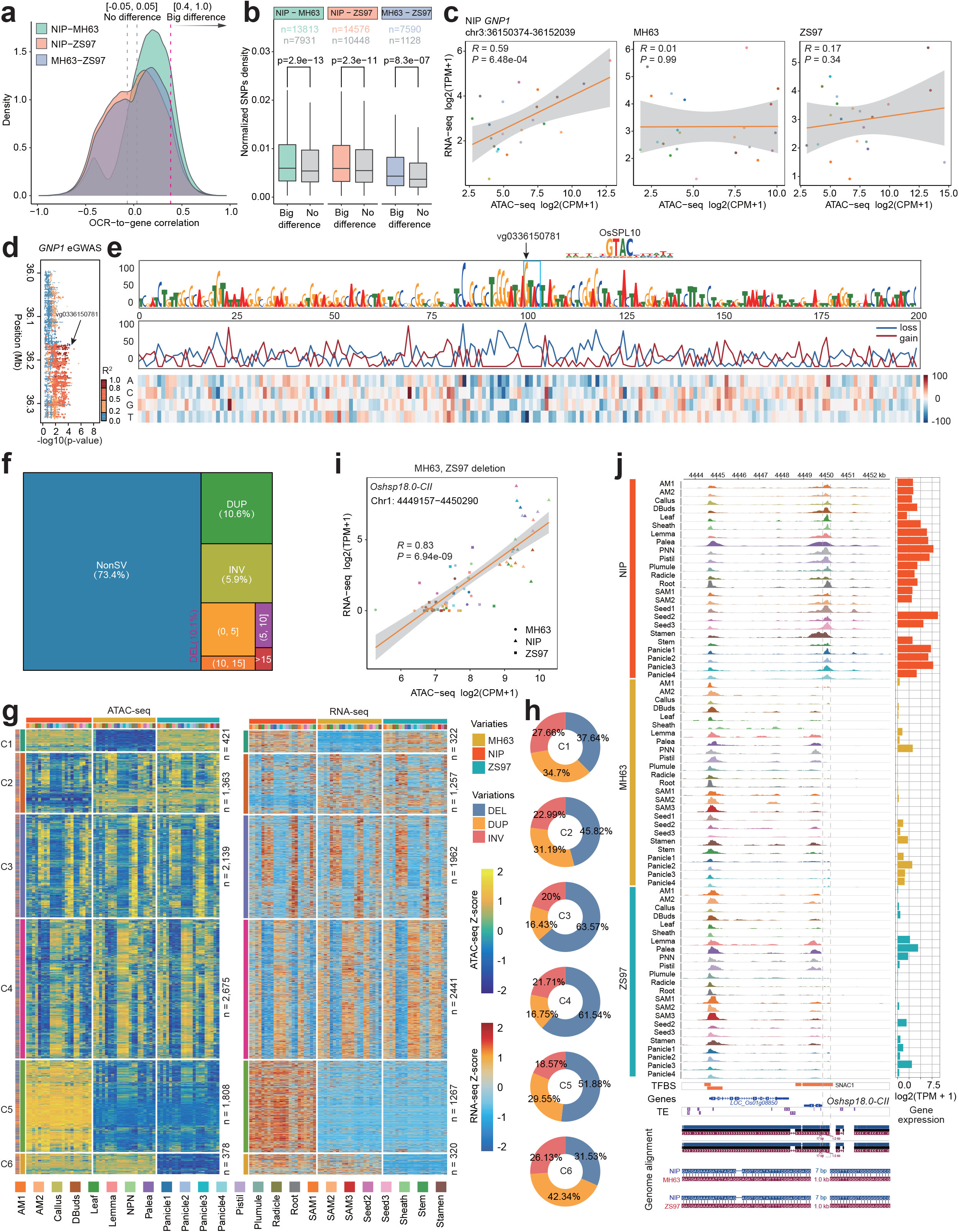
Genomic mutations contribute to cis-regulatory divergence. **a**.Density plot showing the difference in Pearson correlation coefficients (*R*) between the OCR-to-gene of NIP, MH63 and ZS97, respectively. The *R* of OCR-to-gene are not less than 0.4 we consider large differences while *R* located between -0.05 and 0.05 we consider no difference. **b**.Boxplots showing the density of SNP differences between big and small difference groups. Comparisons are made by two-tailed Student’s *t* test. Sample sizes for each group are labeled above their respective boxes. Boxplot shows the median (horizontal line), second to third quartiles (box), and Tukey-style whiskers (beyond the box). **c**.The dot plot demonstrates that the GNP1 gene associates to an OCR (chr3:36150374-36152039) in NIP, but not in MH63 and ZS97 due to the presence of a variant (vg0336150781, G/A). Pearson’s correlation coefficient is used for the test. The error bands indicate 95% confidence intervals. The *P*-values were calculated by two-tailed *Z*-test. **d**.Manhattan plot showing local eGWAS results for *GNP1*. The eGWAS results were obtained from Ming et al^69^. **e**.Changes in chromatin accessibility using deep learning models for mutations of 100 bp each on the left and right of vg0336150781. “Loss” represents reduced chromatin accessibility after the mutation compared to before the mutation, and “gain” represents increased. The figure shows the change in chromatin accessibility before and after the mutation in Panicle2. **f**.The treemap showing the proportion and composition of OCRs without structural variants (SV) and OCRs with SV. Here we only consider deletions (DEL), inversions (INV) and duplications (DUP) for SV. OCRs were considered SV-related when it overlaps with DEL, DUP and INV by at least 1bp. **g**.The heatmap showing the 12,313 OCR-to-gene links (R >= 0.4, *P* < 0.05, two-tailed *Z*-test) associated with SV. They were grouped into 6 clusters based on their chromatin accessibility. The number of OCRs in each cluster and the number of target genes are labeled on the right side of the heatmap. **h**.The doughnut showing the proportion of DEL, DUP and INV in each cluster. **i**.Scatter plot demonstrate Pearson correlation coefficients (*R* = 0.83, *P* < 6.94e-09) between tissues for the accessibility of OCR associated with deletion and the expression of target genes (*Oshsp18.0-CII*). The error bands indicate 95% confidence intervals. The *P*-values were calculated by two-tailed *Z*-test. **j**.Genome Browser showing ATAC-seq signal distribution in the vicinity of gene *Oshsp18.0-CII*. The gray dashed bracket represents the absence of this OCR in MH63 and ZS97 due to the deletion of this sequence. The barplot on the right shows the expression of the gene in each tissue.Source data are provided as a Source Data file.

Overall, the above results provide a valuable resource that can help guide studies of candidate key regulators for tissue-specific gene regulation.

### Systemic localization of GWAS variants in tissue-specific regulatory DNA

Genome-wide association studies (GWAS) have identified numerous natural variations linked to various agronomic traits in rice^3^. To systematically colocalize GWAS-associated variants with the above annotated regulatory elements, especially those from noncoding regulatory regions, we compiled a comprehensive rice GWAS catalog from recent genome-wide association meta-analysis studies^2,53-55^ as well as the NGDC GWAS Atlas database^56^. In total, we collected 4,831 significant (*P* < 1e-5) and representative (only considering lead SNP) associations for 209 distinct quantitative traits which can be classified into seven major categories^57^: morphological characteristics, physiological features, yield components, grain quality, resistance, coloration, and others (Fig. 4a and Supplementary Data 10). In a nutshell, these GWAS SNPs dominantly located in intergenic noncoding regions (Fig. 4b and Supplementary Fig. 8a) and 24.5% of them were either situated within a noncoding OCR (21.1%) or located in linkage disequilibrium (LD) with SNPs in a neighboring OCR (3.4%) (Fig. 4c). Moreover, OCRs revealed significantly higher enrichment of GWAS SNPs than protein-coding sequences (Fig. 4d), highlighting the crucial function of regulatory variants in determining phenotypic characteristics.

Furthermore, our findings demonstrated that OCRs containing GWAS SNPs exhibited greater tissue specificity (Fig. 4e,f and Supplementary Fig. 8b-d). For instance, one of the OCRs containing a GWAS lead variant vg0724671055^54^ (C/T, GWAS *P* < 9.27e-8) significantly associated to panicle number. This OCR was found to be highly accessible specifically to young panicle tissues and its accessibility showed a positive OCR-to-gene link with the expression of *GW7* (*R* = 0.59, *P* < 9.14e-5; Fig. 4g). In another example, the GWAS lead variant vg0431427332 is significantly associated to leaf blade width^53^ (*P* < 1.58e-8), which was located in a SAM/Panicle-specific OCR to positively regulate the expression of *NAL1* (*R* = 0.72, *P* < 1.16e-6) (Fig. 4h). The previous studies have shown that *NAL1* is not only associated with leaf width but also with yield^53^ and has natural variations in expression levels^26^. More examples of validated OCR-related associations are presented in Supplementary Fig. 8e.

### Tissue-specific regulatory variants explain agronomic trait associations

The variation in DNA sequences within OCRs plays a significant role in driving phenotypic innovation through altering chromatin state and gene expression patterns, which usually occurs in a tissue-specific manner. To investigate the relationship between genetic variations associated with agronomic traits and tissue-specific OCRs, we calculated the enrichment of genetic variations within OCRs in a tissue-specific manner. It turned out that significant GWAS SNPs were frequently enriched in OCRs of trait-relevant tissues (Fig. 4f and Supplementary Fig. 8d). For example, GWAS variants associated with spikelet traits were highly enriched in OCRs specific to the tissues of SAM1, pistil and panicle. Motivated by this observation, we performed an enrichment analysis of GWAS-identified SNPs in OCRs from various tissues, using a SNP enrichment method termed CHEERS^58^ (Supplementary Fig. 9). Of the 209 curated GWAS-related traits, ∼78% (163 of 209) phenotypic traits showed GWAS SNP enrichment in at least one tissue (Supplementary Fig. 10 and Supplementary Data 11). The observed enrichment of agronomic trait-associated variants in regulatory elements was highly specific to tissue types, and the association is largely compatible with our current understanding of the tissue function (Fig. 5a). For example, in various GWAS studies, regulatory variants associated with plant height was enriched in stem-related tissues; while genetic associations for grain-related traits (such as grain thickness, grain width, grain length, blighted grains per plant, and filled grains per plant) were highly enriched in OCRs specific to the tissues of seed, lemma, pistil, and stamen (Fig. 5a). Meanwhile, we found that variants associated with root length were predominantly enriched in the root tissue. Specifically, a significant SNP (vg0806201957^59^, *P* < 3.98e-8) located in a root-specific enhancer of *OsHAK12*, which has been shown to be involved in K^+^ uptake in roots^60^ (Supplementary Fig. 11a).

In the case of seed germination percentage, GWAS SNPs were most significantly enriched in plumule-specific OCRs (Fig. 5a). We noted a lead SNP (vg0131729028^61^, A/G, *P* < 8.4e-8) localized within an intronic OCR of *OsbZIP06*, where the intronic OCR and *OsbZIP06* formed a positive OCR-to-gene link (*R* = 0.82, *P* < 2.55e-7) with high tissue specificity in plumule and radicle (Fig. 5b). The minor allele (G) of vg0131729028 was present in a very small proportion (0.3%) in the *XI* population, but in 65.80% of the *Aus* population (Fig. 5c). We mutated the coding region (mainly 1st exon) of *OsbZIP06* with CRISPR/Cas9 and found that the germination rate was higher in two frameshift mutations (osbzip06-1 and osbzip06-2) than in the wild type (Fig. 5d-f and Supplementary Data 12). In contrast, overexpression of the *OsbZIP06* resulted in lower germination rate (Fig.5e,f). Therefore, integration of publish GWAS data and our chromatin landscape can greatly facilitate the identification of candidate genes and the functional annotation of noncoding variants.

Furthermore, when we divided the OCRs into proximal (< 3kb from the TSS, 60,006 OCRs) and distal OCRs (> 3kb from TSS, 35,691 OCRs) before using CHEERS to do enrichment analysis. We observed that the proximal OCRs are more enriched in GWAS SNPs (Fig. 5g-i and Supplementary Fig. 11b). This implies that the enrichment above is mainly driven by the OCR close to the TSS and this result is consistent with previous studies^58,62^.

### Deep learning models accurately predict differences in chromatin accessibility between tissues and unveil common regulatory grammar among varieties

We further investigated whether the tissue- and stage-specific regulatory grammar can be modelled. Deep learning has been successfully utilized to learn and identify essential features in genomic sequences, such as the identification of *cis*-elements^63,64^. Our previous study demonstrated that the Basenji deep learning framework^65^ is powerful for modelling epigenomic data in rice, such as the ability to accurately predict chromatin accessibility and to assess the impacts of variants^7^. Therefore, we optimized the Basenji framework to effectively model our ATAC-seq datasets from multiple tissues (Supplementary Fig. 12a,b). Three distinct models were trained for the varieties of NIP, MH63 and ZS97, demonstrating high accuracy with the mean AUROC values of 0.931, 0.921, and 0.928, respectively (Fig. 6a and Supplementary Fig. 12c). We observed that the Pearson’s correlation coefficient between the predicted and observed values of chromatin accessibility at different locations on the genome reached approximately 0.81, with the best prediction at the location of < 1kb upstream regions (Fig.6b and Supplementary Fig. 12d). This implies that the regulatory syntax patterns within promoter regions could carry more significant information encoded in sequences, which can be effectively captured by deep learning models. Furthermore, the predicted signals from the test sets exhibit the ability to discern between distinct tissues and closely align with the clustering results of the actual values (Fig. 6c). For example, the root-specific expressed gene *RCc3*, responsible for regulating lateral root growth^66^, exhibits distinct chromatin accessibility patterns specifically in root (Fig. 6d and Supplementary Fig. 13).

Subsequently, for each variety-specific model, we used test sets from the remaining two varieties to evaluate the model’s capacity for making predictions across different varieties. Our analysis revealed high Pearson correlation coefficients (about 0.8) between the predicted and observed signals (Fig. 6e). Notably, in the *GSE9* promoter region, there is divergence between *indica* and *japonica* rice, marked by a 9 bp deletion and several SNPs in MH63 when compared to the sequences of NIP and ZS97^67^. The ZS97 model predicted the chromatin accessibility of this region in MH63 with weak signals. Contrarily, the ZS97 model accurately predicted the chromatin accessibility in NIP and ZS97, showing strong signals (Fig.6f and Supplementary Fig. 14). These results suggest that the deep learning model can effectively make accurate predictions across varieties, implying that shared regulatory grammar across rice varieties.

We next performed comparative analyses on ATAC-seq data of 22 matched tissues/organs in both *japonica* rice (NIP) and *indica* rice (MH63 and ZS97), utilizing their respective reference genomes (Fig. 1a,b). We found that roughly 60% (60,764 out of 95,697) of OCRs were shared across all three cultivars (Fig. 6g and Supplementary Data 13). The *indica* varieties MH63 and ZS97 exhibited a higher proportion of shared OCRs compared to NIP which from different subspecies (Fig. 6g). We next sought to compare chromatin accessibility dynamics of the 1:1:1 orthologous OCRs across the three varieties (referred to as triads; see *Methods*). To investigate the accessible bias of orthologous OCRs, we compared the chromatin accessibility of orthologous OCRs in each individual tissue (Fig. 6h). Orthologous OCRs were assigned into seven categories on the ternary plot based on their relative accessibility, including a balanced category and six dominated or suppressed categories in specific cultivars (Supplementary Fig. 15). The proportion of OCR triads assigned to unbalanced categories varied among different tissues, ranging from 3.2% in plumule to 24.8% in AM1 (Fig. 6h and Supplementary Fig. 16a). While promoters generally display balanced OCRs, indicating consistent accessibility across different cultivars, enhancers frequently exhibit unbalanced OCRs, reflecting cultivar-specific regulation (Supplementary Fig. 16b). Interestingly, unbalanced OCRs harbored more genotypic variations in terms of SNPs (Fig. 6i). This observation led us to suppose whether sequence variation among different varieties caused the differences in chromatin accessibility of these OCR orthologs. Therefore, we used NIP-based deep learning model to predict the chromatin accessibility signals of sequences from orthologous OCRs in NIP, MH63 and ZS97, respectively, and then compare these predictions. The results showed that about 50% of the differences in orthologous OCRs could be successfully resolved in terms of sequence variation (Fig. 6j and Supplementary Fig. 17).

In summary, the above results illustrate that deep learning models could accurately predict chromatin accessibility across tissues and varieties. The high accuracy of the models also indicates the high quality of our data.

### Elucidate key genetic changes underlying cis-regulatory divergence by deep learning models

Genetic variants and *de novo* mutations in regulatory regions may lead to *cis*-regulatory divergence and thus changes in gene expression and organismal phenotypes^68^. We systematically dissected the *cis*-regulatory divergence due to genomic sequence changes (e.g., SNPs) in regulatory regions, which could be inferred from ATAC-seq data. To measure the effect of the variant on chromatin accessibility, we extracted variants that differed in the three varieties. The effect of different alleles of each variant on chromatin accessibility was evaluated using the deep learning models. We found that unbalanced OCRs had a higher absolute effect score than the balanced OCRs (Supplementary Fig. 18a) and these large-effect loci were significantly enriched for eQTLs^26,69^ (Supplementary Fig. 18b). This observation suggests that these putative large effect variants are associated with changes in chromatin accessibility and gene expression. Meanwhile, we performed separate OCR-to-gene correlation analysis for each of the three varieties. We then identified conserved OCR-to-gene links and compared the correlation coefficients between them (Fig. 7a). Notably, OCRs with significant differences in correlation coefficients exhibited higher SNP density (Fig. 7b), and the OCR-to-gene links with large differences in correlation coefficients between MH63 and ZS97 were significantly enriched for differential *cis*-eQTL between MH63 and ZS97 (Fisher’s exact test, odds ratio = 1.81 and *P* < 1.83e-28)^70^. These suggesting that regulatory sequence variations among different varieties could influence gene expression. For instance, we observed that a SNP (vg0336150781, G/A) located in the *GNP1* promoter region control grain number and plant height^71^. Among the OCR-to-gene links we inferred, the allele in NIP (‘G’ at this SNP) correlated with *GNP1* (*R* = 0.59, *P* < 6.48e-04), whereas the allele (‘A’ at this SNP) did not show OCR-to-gene correlation in MH63 (*R* = 0.01, *P* = 0.99) and ZS97 (*R* = 0.17, *P* = 0.34) (Fig. 7c). In addition, eGWAS also demonstrated that this SNP affects *GNP1* expression (Fig. 7d). When we evaluated the effects of this SNP with the deep learning model, we found that mutation of this SNP from “G” to “A” in Panicle2 significantly reduced chromatin accessibility (Fig. 7e). We also found that this variant overlaps with the footprint of OsSPL10 identified in Panicle2, and its binding site shows the typical “GTAC” motif of the SBP TF family. These results suggest that mutations control gene expression by affecting TF binding to alter chromatin accessibility.

Besides point mutations, small genomic alterations (including short insertions/deletions, inversions, and duplications) may abolish OCRs and thus confer an important avenue of regulatory divergence. We quantified all OCRs based on the NIP reference genome and investigated whether their regulatory activity dynamics were associated with short genomic alterations, which were determined by whole genome comparison across different cultivars (see *Methods*). In total, we found that nearly one third (26.6%) of the OCRs harbored small alterations (Fig. 7f). The regulatory activity of these mutation-associated OCRs is positively correlated with their surrounding gene expression patterns in a cultivar-specific manner (Fig. 7g,h), as exemplified at the loci of *Oshsp18*.*0-CII* and *MAG2* (Fig. 7i,j and Supplementary Fig. 19a). Notably, GO analysis showed that these genes were highly enriched for various ‘response’ related functions (Supplementary Fig. 19b and Supplementary Data 14). Further investigation revealed that the identified mutation-embedded OCRs were significantly overlapped with transposable elements (TEs) (Supplementary Fig. 19c). The above results indicate that TEs may contribute to modification of regulatory sequences, fine-tuning gene expression networks and driving new functions^72^.

## Discussion

Despite substantial progress, a complete catalog of regulatory sequences within the rice genome remains elusive, limiting the understanding of tissue-specific regulatory dynamics and GRNs. Our study presents a comprehensive exploration of rice genome regulation using the UMI-ATAC-seq technique^12^, providing insights into tissue-specific regulatory elements and their influence on complex agronomic traits. Of note, the identified OCRs in rice encompass approximately 15% of the genome, a notably higher proportion compared to previous reports in plants such as Arabidopsis (∼4%)^73^ and maize (∼4%)^74^. This expanded coverage underscores the importance of sampling depth in characterizing the regulatory complexity in plants and highlights the need for further comparative analyses to elucidate species-specific regulatory features.

Predicting OCR-to-gene links presents a significant challenge due to the intricate regulatory mechanisms governing gene expression. By integrating RNA-seq data from matched tissues, we predicted 59,075 OCR-to-gene links, including many reported enhancer-to-gene links. This analysis offers a holistic view of how changes in chromatin accessibility directly impact gene expression patterns, underscoring the significance of regulatory elements in shaping the rice transcriptome. The identified associations between enhancers and target genes provide guidelines for dissecting complex regulatory mechanisms and gene editing in non-coding regions. The approach for predicting OCR-to-gene links based on multi-omics data is versatile and transferable to other plant species. Despite our efforts to predict OCR-to-gene links, less than half of the protein-coding genes exhibit a relatively strong correlation (Pearson correlation coefficient |*R*| ≥ 0.4 and *P* < 0.05) with OCRs. The complexities of dynamic and context-dependent regulation, coupled with long-range and indirect regulatory mechanisms, introduce additional layers of complexity to OCR-to-gene link prediction beyond the capabilities of linear models aimed at directly mapping OCRs to their target genes. These factors likely contribute to the weaker correlations observed for certain genes. Moreover, tissue-specific and housekeeping genes are difficult to correlate through linear models due to the small variation in expression levels between tissues (Supplementary Fig.20).

Deep learning has emerged as a potent tool for interpreting the genomic and epigenomic data^63,64^, but its application in rice is hindered by the scarcity of high-quality epigenomic datasets. Our study addressed this gap and successfully modelled the chromatin accessibility of three rice varieties. The highly accurate models enable the prediction of chromatin accessibility variation across varieties using sequences, providing a reference for scientists to explore the functional effects of rare variants or new variants across different tissues.

Moreover, our comparative analysis across varieties revealed *cis*-regulatory divergence that could largely be predicted using deep learning models based on sequences, highlighting the genetic diversity of rice varieties and its impact on regulatory architecture. By integrating GWAS data, we localized significant variants within noncoding regulatory regions, demonstrating that these variants are preferentially located in tissue-specific OCRs, thus providing insights into the influence of regulatory variations on phenotypic outcomes. A notable achievement of our study is the identification of OsbZIP06’s role in seed germination, demonstrating the potential of integrating GWAS data with chromatin accessibility to uncover the genetic basis of complex traits.

In summary, our extensive chromatin accessibility atlas and the deep learning models we have constructed not only enhance our understanding of regulatory elements in rice but also serves as a versatile resource for gene editing and breeding strategies targeting non-coding regions. Nevertheless, there are several limitations associated with our study. Firstly, our map solely encompasses data from normal conditions, omitting insights into responses to biotic or abiotic stresses, mutants, and diverse environmental circumstances. Secondly, the inferred associations between OCRs and genes require experimental validation to confirm their regulatory relationships. Thirdly, our study primarily employed the NIP reference genome, thereby excluding sequences that were not available in the NIP genome. Furthermore, the advent of single-cell technologies has opened avenues for studying *cis*-elements at a single-cell resolution^74,75^. In the future, incorporating single-cell data will be crucial for further characterizing the heterogeneity among different cell types. These advancements will collectively contribute to a more comprehensive understanding of the regulatory landscape in rice and beyond.

## Methods

### Plant materials, ATAC-seq, and RNA-seq experiments

Three rice varieties, Nipponbare, Zhenshan 97 and Minghui 63, were planted in a field in Wuhan, China in the summer of 2020 and were used to obtain most of the tissues or organs used in this study. Details of the sampling are listed in Supplementary Data 1. We followed our previously established method to perform UMI-ATAC-seq experiments^12^. RNA was isolated using TRIzol reagent (Invitrogen Life Technologies), and sequencing libraries were prepared using the MGIEasy RNA Library Preparation Kit. The libraries were subsequently sequenced on the MGISEQ-2000.

### ATAC-seq data analysis

For the pre-processing of ATAC-seq data, we follow the workflow of ChIP-Hub^14^ and cisDynet^76^. The raw reads were first trimmed by Trimmomatic (v.0.36)^77^ to remove sequencing adapters. The trimmed reads were aligned to the *Oryza sativa L*.*ssp*.*japonica* (cv. Nipponbare) reference genome (v.7.0)^16^ using Bowtie2^78^ with the following parameters “-q—no-unal—threads 8—sensitive”. All reads mapped to mitochondrial and chloroplast DNA were removed. After sorting mapped reads with SAMtools^79^ (version 0.1.19), we only used properly paired reads with high mapping quality (MAPQ score > 30) for the subsequent analysis. The PCR duplicates were removed using the MarkDuplicates function from Picard tools (version 2.60; http://broadinstitute.github.io/picard/). The “callpeak” function in MACS2^80^ (version 2.1.0) was used to call peaks with the following parameters: “-g 3.0e8 --nomodel --keep-dup 1 -B --SPMR --call-summits”. The “-shift” used in the model was determined by the analysis of cross-correlation scores using the phantompeakqualtools package (https://code.google.com/archive/p/phantompeakqualtools/).

### RNA-seq data analysis

RNA-seq reads were aligned to the Nipponbare reference genome^16^ using STAR^81^ (version 2.7.1a). The expression of annotated genes was measured by RSEM^82^ (version 1.2.22) and normalized with transcripts per million (TPM).

### Linking OCRs to target genes

To assign OCRs to genes, we used an approach similar to the previous study^23,83^. First, we prepared the ATAC-seq quantification matrix, with each row representing a merged OCR and each column representing a sample. After merging replicates, 66 tissues with both ATAC-seq data and RNA-seq data were taken as independent samples for the analysis. For the gene expression quantification matrix, we removed possible noise by considering only those genes whose TPM of each row added up to > 1.5. For each of the remaining 29,571 genes, we screened the OCRs that might regulate the gene within 20 kb upstream and downstream of the TSS of that gene separately. Then we calculated the Pearson correlation coefficients between the chromatin accessibility of these OCRs and the expression of that gene. Then we randomly generated pseudo-peak sets of the same length and number as these OCRs on other chromosomes, repeated the process 10,000 times, and used *Z*-test (*z*.*test* function from the *R* package ‘TeachDemos’) to calculate *P* values. Finally, we considered that absolute Pearson correlation coefficients (|*R*|) >= 0.4, and *P* < 0.05 were significant OCR-to-gene links. For the identification of OCR-to-gene links of NIP, MH63, and ZS97, we used the same strategy except that we used ATAC-seq and RNA-seq samples of the corresponding varieties.

### Tissue-specific OCRs analysis

We merged the peaks with NIP tissues, counted the number of Tn5 cuts of these peaks in different tissues, normalized them and then used the Jensen-Shannon Divergence (JSD) from the philentropy *R* package (https://github.com/drostlab/philentropy) to screen tissue-specific OCRs. Here we considered OCRs with JSD score > 0.08 (except > 0.1 for young panicle) as tissue-specific OCRs. For tissue-specific OCR, we performed Z-score transformation by row for visualization. To identify the top tissue-specific OCRs in each tissue, we employed a scoring metric known as Specificity Measurement (SPM), as detailed in the method provided at https://github.com/apcamargo/tspex. Subsequently, we sorted the OCRs within each tissue based on their SPM scores to select the top 2500 tissue-specific OCRs in each tissue.

### Motif enrichment analysis

For motif enrichment analysis of tissue-specific OCRs, we first calculated the Tau index score using SPM metric for each tissue’s OCRs and selected the top 2,500 OCRs of each tissue for motif enrichment analysis according to the ranking of Tau index scores. Then we used GimmeMotifs^37^ with maelstrom function to determine the tissue-specific motifs enrichment. We set the “--filter-cutoff” to 0.4. The input Position weight matrix (PWM) was downloaded from the JASPAR^84^ database (https://jaspar.genereg.net/). We combined the enrichment result of three methods (Lasso, Bayesian ridge regression, and boosted trees regression) to get the final motif enrichment lists.

### ChIP-seq enrichment analysis

The public ChIP-seq data used in this study are provided in Supplementary Data 3. We downloaded the narrow Peak files of these TFs from the ChIP-Hub database (https://biobigdata.nju.edu.cn/ChIPHub/), and then used BEDTools^85^ (version 2.29.1) fisher function to calculate the enrichment level with OCRs.

### TF motif and footprinting analysis

For the TF motif enrichment analysis, we used the SEA program from the MEME suite and used constitutive OCRs as background. We considered motifs with *P*-value < 1e-5 to be significantly enriched. For genome-wide TF potential binding sites, we used the FIMO program in MEME to identify them and also used *P*-value < 1e-5 as the cutoff.

TF footprints were calculated by TOBIAS (version 0.13.1)^86^. We first used TOBIAS ATACCorrect function to correct the Tn5 inherent insertion bias. Then we calculated the footprint score in OCRs using FootprintScores function with default parameters. Finally, we used BINDetect function to predict transcription factor binding footprint for each sample, which were matched to curated list of JASPAR^84^ motifs (https://jaspar.genereg.net/).

### Cross-variety comparisons of OCRs

We first aligned the whole genome sequences of NIP, MH63, and ZS97 to each other. The strategy used for the whole-genome alignment was similar to the previously described method ^23^. The results were further filtered to obtain more reliable conserved sequences following the default process of “Reciprocal Best” (http://genomewiki.ucsc.edu/index.php/HowTo:_Syntenic_Net_or_Reciprocal_Best). We obtained three superset OCRs by merging OCRs of tissues shared by three varieties s (n = 22). Then we used the bnMapper.py script in bx-python (https://github.com/bxlab/bx-python) to convert the OCRs coordinates of MH63, ZS97 to the corresponding coordinates of the NIP. We then considered the OCRs of MH63, ZS97 with at least a 50% overlap with the OCRs of NIP to be conserved OCRs for the three varieties. To obtain a quantitative matrix of conserved OCRs, we first quantified all OCRs for each variety and divided the length of the corresponding OCRs by the CPM strategy, and then extracted the conserved OCRs for each variety for subsequent analysis. We then refer to it to classify conservative OCRs into seven categories (NIP dominant, MH63 dominant, ZS97 dominant, NIP suppressed, MH63 suppressed, ZS97 suppressed, and balanced).

### Deep learning model analysis

We used the Basenji^65^ deep learning framework with modifications to accommodate the relatively small rice genome for deep learning model training. We first use the *bam_cov*.*py* script to convert the bam files into bigwig files. We then used the *basenji_data*.*py* script to prepare the input files for the deep learning model according to the following parameters: “-d 1.0 -s 0.1 –local -t 1 -v 4”. The “-c, -l, -w” of these parameters are shown in the Supplementary Data 15. Data from chromosome 1 was used as the test sets and data from chromosome 2 was used as the validation sets. Next we used the basenji_train.py script to train the model on a NVIDIA GTX 3090. The *basenji_test*.*py* script (default parameters) was used to perform the model performance test. We found differences in the training performance for different parameter settings for rice, with “-l 32768 -c 2048 -w 128” being the best, and subsequent analyses were based on models trained with this parameter. To measure the effect of variation in OCRs, we used the *basenji_sat_bed*.*py* script to perform base mutations at this locus and calculated the difference in signals between the reference and the mutation as the variation effect value. To predict the chromatin accessibility of orthologous OCRs, we extended the centre of the OCR by 16,384 bp left and right to make a total length of 32,768 bp. Sequences exceeding the length of the corresponding chromosome were removed, and then sequences of the corresponding varieties were extracted using BEDTools getfasta function, and then *basenji-predict_bed*.*py* was modified to enable it to use fasta as input.

### Analysis of structural variants and transposable elements

We downloaded deletions, duplications and inversions for MH63 (CX145), ZS97 (B156) in Rice SNP-Seek Database (https://snp-seek.irri.org/). Since this database provides large structural variants with a minimum length of 10 bp, we also integrated a series of variants with reference to this workflow^7^. Briefly, we selected Leaf ATAC-seq data from NIP, MH63 and ZS97 varieties, then aligned them to the Nipponbare reference genome using BWA-MEM^79^ (version 0.7.12-r1039) and identified INDELs using GATK^87^ (version 3.3-0-g37228af). The annotation files of transposable elements (TEs) were downloaded from Phytozome database. We use the default parameters of the BEDtools^85^ (version 2.29.1) intersect function to identify OCRs that overlap with structural variants and TEs.

### GO enrichment analysis

All GO enrichment analysis was done in the Rice Gene Index database (https://riceome.hzau.edu.cn/)^88^ using default parameters. We considered FDR < 0.05 as a significantly enriched pathway.

### GWAS data processing

The genotype and phenotype data used in this study were downloaded from four published cohorts. We refer to this reference^3^ to name them as 529 rice accessions^2^, 1,275 Chinese rice accessions^54^, 176 Japanese rice accessions^53^ and 3K rice accessions^55^, respectively. GWAS was performed separately for each cohort by GCTA^89^ (version 7.93.2) with mixed linear model. To determine the significant SNPs cutoff, we first used Genetic type 1 error calculator (GEC^90^, version 0.2) to evaluate the effective numbers of independent SNPs (*N*) and approximated by 0.05/*N* to estimate the cutoff. The threshold for significant SNPs varied by cohorts, we set the thresholds to 1⁳×⁳10^™6^, 1⁳×⁳10^™4^, 1⁳×⁳10^™5^, and 1⁳×⁳10^™6^ for 3K rice accessions, 176 Japanese rice accessions, 529 rice accessions, and 1,275 Chinese rice accessions, respectively. Variants with a minor allele frequency (MAF) that was < 5% were excluded. For the lead SNP identification, we used PLINK^91^ (version 1.9) and set the parameter “--clump-p1” to the threshold we defined above, “--clump-p2 0.05 --clump-r2 0.6 --clump-kb 1000” for the first round of parameters. Then we set the second round of “--clump-r2” to 0.1, other parameters are unchanged. We used PLINK with the following parameters “--ld-window-kb 1000 --ld-window 99999 --ld-window-r2 0.8” to calculate the SNPs with strong linkage disequilibrium (*r*^2^ >0.8) with lead SNPs.

### Enrichment analysis of GWAS-associated SNPs of different *P*-values with OCRs

The enrichment in the OCRs of a tissue at a given threshold of different *P*-values was calculated as the fraction of SNPs with *P*-values below this threshold that overlap with the OCRs (merged all NIP tissues’ OCRs), divided by the fraction of all noncoding SNPs that overlap with the OCRs in the study. Enrichment was performed at *P*-value thresholds ranging from 1e-1 to 1e-7. The smallest threshold had at least 50 SNPs in their study to ensure the sufficient sample size.

### GWAS SNPs enrichments

We first merged the peaks from all tissues in Nipponbare and used this peak superset to quantify each tissue. To make sure that our analysis was not interfered by low confident peaks, we dropped the peaks in the tenth percentile of the lowest Tn5 cuts coverage, yielding 86,011 ATAC-seq peaks finally. Then we performed the normalization with *CHEERS_normalize*.*py* from CHEERS^58^ (Chromatin Element Enrichment Ranking by Specificity) software (https://github.com/TrynkaLab/CHEERS/tree/python3). The normalized quantification matrix was next transformed to tissue-specificity score with range 0-1. To do the enrichment analysis, we used the set of lead SNPs and SNPs with strong linkage disequilibrium (*r*^2^ > 0.8) with the lead SNPs computed separately for the corresponding cohort from the 209 GWAS above as the input to *CHEERS_computeEnrichment*.*py*. The enrichment *P*-values were transformed by -log10 and normalized by row with Z-score for visualization. For the proximal and distal GWAS SNPs enrichment analysis, we first divided OCRs into proximal and distal according to its summit distance from the nearest TSS. All other steps are the same as described above.

### Generation of transgenic rice plants

To obtain overexpression lines of *OsbZIP06*, the cDNA of *OsbZIP06* was cloned using primers OsbZIP06-OE-F and OsbZIP06-OE-R and inserted into the Kpn1-BamH1 site of the pCAMBIA1301 vector and fused with the maize Ubiquitin promoter and three FLAG tags at its C-terminus using the ClonExpress II One Step Cloning Kit (Vazyme). The construct was then transformed into ZhongHua11 (ZH11) by Biogle GeneTech. Primers used to clone *OsbZIP06* are listed in Supplementary Data 16.

For the *OsbZIP06* mutant strain, T1 generation seeds produced using the CRISPR-Cas9 system were purchased from Biogle GeneTech. The sgRNA sequence OsbZIP06-CR-gRNA is listed in Supplementary Data 16.

### Seed germination experiments

Seed germination experiments were performed as previously described^61^. Seeds of Zhonghua 11, OsbZIP06 mutant in the Zhonghua 11 background were used for germination experiments.

### Statistics and reproducibility

If not specified, all statistical analyses and data visualization were done in *R* (version 4.0.0) or Python (version 3.8.9). *R* packages (e.g. ggplot2 and plotly) and Python packages (e.g. Seaborn) are heavily used for graphics. All the sources data for each figure can be found in the Supplementary Information. Specific tests used to determine statistical analyses are noted in each figure legend.

## Data Availability

The sequencing data from ATAC-seq and RNA-seq generated in this study have been deposited in the NCBI BioProject database under accession code PRJNA940508 [https://www.ncbi.nlm.nih.gov/bioproject/PRJNA940508]. All public ChIP-seq used in this study are download from ChIP-Hub database (https://biobigdata.nju.edu.cn/ChIPHub/). The accession number are provided in the Supplementary Data 3. Some critical analysis results about this study can be accessed in the CART database (https://biobigdata.nju.edu.cn/cart/). Source data are provided with this paper.

## Code Availability

The code related to figures is available at https://github.com/compbioNJU/CART.

## Acknowledgements

This work is supported by grants from STI2030-Major Projects (2023ZD04076), the National Natural Science Foundation of China (No. 32070656, 31922065), the Hubei Provincial Natural Science Foundation of China (2023AFA043), the Earmarked Fund for the China Agriculture Research System (CARS-01-01), the Fundamental Research Funds for the Central Universities (2662023PY002) and HZAU-AGIS Cooperation Fund (SZYJY2023003). The authors acknowledge the Center for Information Technology and the High-Performance Computing Center of Nanjing University and the bioinformatics computing platform of the National Key Laboratory of Crop Genetic Improvement at Huazhong Agricultural University for providing high performance computing (HPC) resources.

## Author contributions

D.C. and W.X. conceived and designed the project. T.Z., C.X., X.X., Y.L., J.Y. performed experiments. T.Z., D.C., X.Z., R.Y., L.W., Z.Z., L.M. and Y.Y. conducted the bioinformatics analysis. D.C., W.X and T.Z. wrote the paper. All the authors reviewed and approved the paper.

